# Stochastic bounds of aggregation dynamics distinguish near-wild-type from wild-type strains in social bacteria

**DOI:** 10.1101/2023.02.26.530117

**Authors:** Merrill E. Asp, Eduardo A. Caro, Roy D. Welch, Alison E. Patteson

## Abstract

The genotype-to-phenotype problem (G2P) for multicellular development asks how genetic inputs control collective phenotypic outputs. It is a difficult problem even to observe. On the genotype side, the phenotypic impact of mutation is often subtle due at least partly to gene redundancy and myriad other factors. On the phenotype side, biological and even technical developmental replicates can display significant phenotypic variation due at least in part to stochasticity, again with other factors. We attempt to partially resolve the G2P inputs and outputs from the obfuscating effects of factors like redundancy and stochasticity. As a model organism, we selected the biofilm-forming species *Myxococcus xanthus*, a motile self-organizing bacterium that forms three-dimensional cell aggregates that grow and mature into spore-filled fruiting bodies when under starvation stress. We developed data acquisition tools and analysis and visualization methods that can produce a topological map of *M. xanthus* development. We demonstrate that even subtle effects on developmental dynamics caused by mutation can be identified, discriminated, characterized, and given statistical significance.

## Introduction

Development is an energetically expensive and complicated part of many organisms’ life cycles. For a genetically specified multicellular phenotype to incarnate, the process requires the stepwise self-organization of increasingly ordered states and an active and robust dampening response to disruptive forces, such as environmental stress and mutation^1–3^. The control systems for development involve non-linear redundant branching and intersecting intracellular and intercellular signal transduction pathways that provide spatiotemporal coordination of the transcriptional, translational, and post-translational events required for the organism to manifest^4,5^. This complicated genotype-to-phenotype problem, abbreviated as G2P, is the broad task of understanding the iterative process of genetic cause and phenotypic effect that eventually results in biological emergence.

We use the Gram-negative delta-proteobacterium *Myxococcus xanthus* as a model organism. In the laboratory on an agar substrate, *M. xanthus* exists as a single-species motile biofilm called a swarm. Under nutrient-rich conditions, a swarm will expand across an agar surface as its component cells grow and move (swarming). In contrast, under non-nutritive (starvation) conditions, a swarm will not expand, although its component cells are in fact moving around faster within it. Instead, a starving swarm will undergo a transformation over a period of approximately one day, during which swarm cells organize into a discrete number of mound-shaped aggregates distributed non-randomly across the swarm area, with each aggregate harboring a bolus of thousands of cells. Over the next few days, the cells at the center of each aggregate differentiate into quiescent myxospores, at which point the aggregates are considered to have matured into a fruiting body. The entire process represents a rudimentary but robust form of multicellular development and can therefore be studied as an example of G2P^6^.

Development of a multicellular prokaryote, such as the formation of an *M. xanthus* fruiting body, may be less complicated than development of a eukaryote, such as the formation of a mouse, fly, or flatworm, but it is still complicated enough to involve hundreds of genes arranged in branching networks of intersecting pathways. Many of the genes known to be involved in these networks and pathways were first identified through mutation and phenotypic characterization^7^. It is a foundational protocol in developmental biology: first, a wild-type (WT) strain is selected and its development phenotype is characterized, then mutations are introduced into the WT genome to create new (mutant) strains, and their development phenotypes are characterized, then the phenotypes of the mutant strains are compared to WT and, if a strain displays a significant deviation, the gene(s) and other genetic element(s) affected by the mutation are deemed more likely to be involved in development. This information can then be used for genome annotation and to guide future research.

Permutations of this protocol have been used many times on *M. xanthus* to provide preliminary biological process annotations for a significant percent of the genes in its genome. These annotations are more impactful when they can be used for the purpose of comparison and, to be objectively comparable, their underlying phenotypic data must be quantitative. A mutant strain’s descriptive (qualitative) phenotype characterization can be used for comparison to WT only if its deviation is catastrophic, in which case mimics being quantitative because the possible outcomes are binary (success/failure). Phenotypic deviations from WT that are less than catastrophic are still potentially quantitative, if measurable features (dimensions) can be identified that match the descriptive characterization, or at least aspects of it. In fact, if the quantification of features is reproducible and statistically significant, they can be used to differentiate mutant strains from WT, even when their descriptive characterizations are nearly identical.

A requisite condition to establish significance when characterizing and quantifying a developmental phenotype is a defined boundary that distinguishes the phenotype of WT from a near-WT mutant. There are at least three confounding factors that make this difficult. First, developing biological systems exhibit an inherent phenotypic stochasticity, and efforts at holding genome sequence and experimental conditions constant can only reduce the variation to a nontrivial baseline. Second, biological systems are also phenotypically robust to the impact of mutation because evolution guides the genes, networks, and pathways that control development to incorporate redundancies due, at least in part, to mechanisms like duplication and divergences^8^. Third, developmental processes are often difficult to observe, record, and analyze in replicates sufficient to establish significance.

Statistical techniques that are useful in the face of highly stochastic events such as gene expression are a topic of active interdisciplinary research^9^ and the inherent ability of living systems to submit to statistical study is an open epistemological question^10^. From a strictly statistical perspective, however, if a baseline level of variation can be determined for WT, any differences distinct from that baseline are capable of distinguishing phenotypes. At present it is an interesting idea that must be verified through experiment.

In this study, we describe the design, construction, and operation of an experimental setup for observing many instances *M. xanthus* development and an analysis pipeline that quantifies different features of the development phenotype and displays them as a map. We then employ this setup to distinguish between WT *M. xanthus* development and four near-WT mutants.

**Figure 1:**
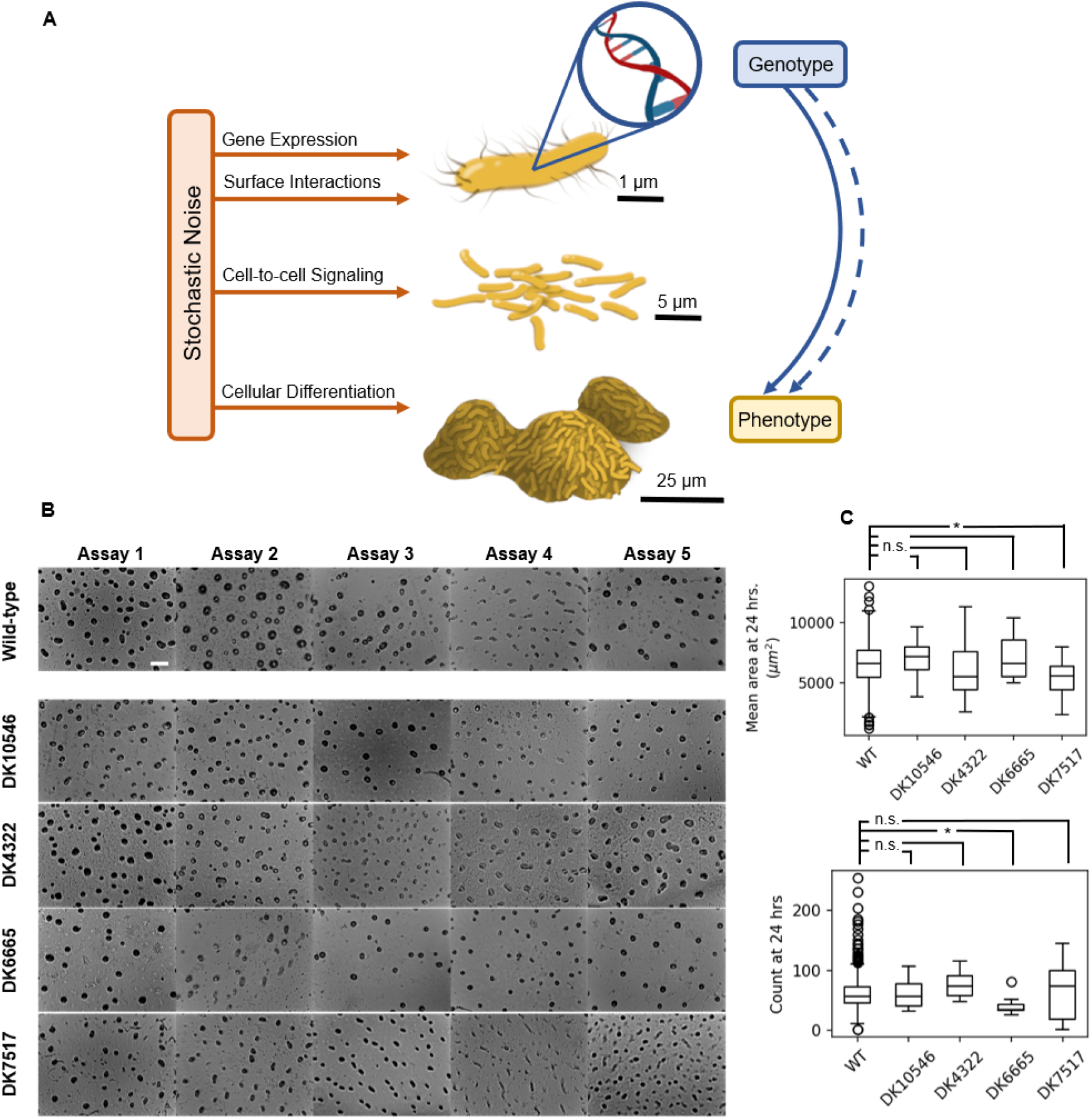
Stochasticity is inherent to multicellular behaviors in social bacteria. (**A**) A bacterial colony undergoing fruiting body development is exposed to stochastic noise on multiple scales. At the cellular level, gene expression depends on thermally driven chemical events, and environmental factors such as variations in temperature and humidity introduce further uncertainty. Thus both direct and indirect effects of genotype arrive at a final phenotype, possibly via multiple developmental paths. (**B)** Images of final developmental phenotype for separate aggregation assays at 24 hours post inoculation. Pictured are a range of outcomes from the wild-type *M. xanthus* strain as well as the four mutant strains used in this study. Aggregates are visible as dark spots, seen from above. Scale bar, 250 μm. **(C)** The average final area and final count of wild-type aggregates and those for four mutant strains are reported with boxplots. Although there are some differences in these typical metrics of comparison, there is considerable overlap between wild-type and each of the mutant strains. N > 500 measurements for wild-type, taken over 25 different days; N = 15 measurements for each mutant strain, taken over 2 different days.

## Results

An accurate measurement of the variation of wild-type behavior requires a large sample size to accommodate the many developmental outcomes observed in *M. xanthus* fruiting body development and estimate their probability. To accomplish this with enough simultaneously running assays to measure the impact of day-to-day variation, we produced custom built microscopes and a microscope control network to collect and organize developmental data. Each fruiting body aggregation assay requires a sealed chamber with sufficient temperature, oxygenation, and humidity for development to occur. Cells are inoculated from liquid culture and sealed in each slide assembly, where the aggregates begin to form after about five hours. After manual setup and focus, automated imaging proceeds for 24 hours, at which point most aggregates are stable. The resulting time series images are organized on a central hub computer from which image processing can begin.

**Figure 2:**
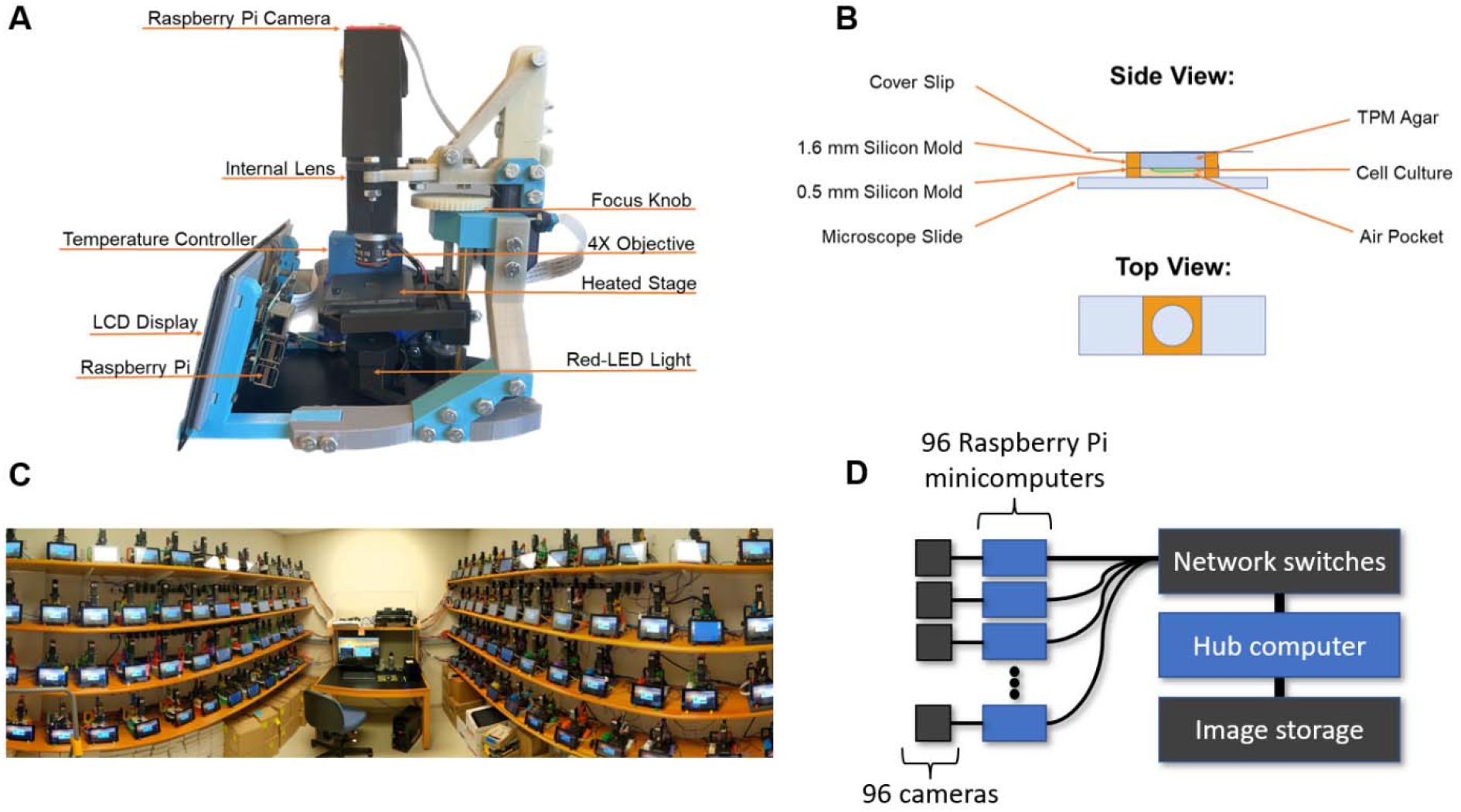
High-throughput time series acquisition setup. **(A)** A single microscope, at 1.2 kg and 24×19×25 cm^3^, with 3D-printed armature, 4X objective lens, light source, heated stage, camera, and Raspberry Pi microcomputer. **(B)** Slide assembly for each developmental experiment. Sandwiched between a glass coverslip and a glass slide, two silicone gaskets create a sealed enclosure containing a disk of non-nutritive agarose on which a colony of *M. xanthus* has been inoculated. Aggregate development is imaged over a 24-hour period with one image taken each minute. **(C)** Panoramic photograph of full image acquisition setup including 96 microscopes and central hub computer. **(D)** Basic network architecture for centralized image storage and control of all 96 microscopes.

Using a dataset of over 500 wild-type aggregation time series acquired over 25 separate days, we quantify the range of developmental phenotype by measuring ten quantitative metrics for each video. We choose three metrics related to timing: start time, when aggregation begins; peak time, when the area occupied by aggregates reaches its maximum; and stability time, when the number of aggregates becomes stable. We also measure the mean and standard deviation in average aggregate area at two time points: peak time, and 24 hours. We measure the number of identifiable aggregates at both peak time and 24 hours. Finally, we measure the fraction of aggregates that appear and then disperse before 24 hours elapse from inoculation.

Myxobacteria development, when observing fruiting body morphogenesis, is typically resolved by 24 hours, especially in the wild-type strain DK1622. After placement onto starvation media, the movement of aggregates, their size, and distinctive morphology has usually been set by this point, even though the internal myxospores are still maturing for an additional 3-5 days. For the purposes of our study, we chose to limit our time lapses to 24 hours to highlight differences in the dynamics of fruiting body aggregation. Phenotypic differences could yet be observed in mutants that take longer than 24 hours using the methods established here. The specific algorithms used to determine each phenotypic metric are detailed in the Supplementary Materials (Table S2). These metrics are extracted with a custom Python image processing algorithm that identifies and measures each aggregate, as described in Methods. Values for these metrics across the wild-type dataset are shown in Figure 3A, with the distributions illustrating averages and variation for each metric. Peak time, aggregate count at peak time, and the fraction of aggregates that disperse exhibit bimodal distributions. Long-tailed distributions, such as the standard deviation (σ) of area and aggregate count (both at peak time and after 24 hours) indicate the presence of abnormal phenotypes with extreme values in these metrics.

**Figure 3:**
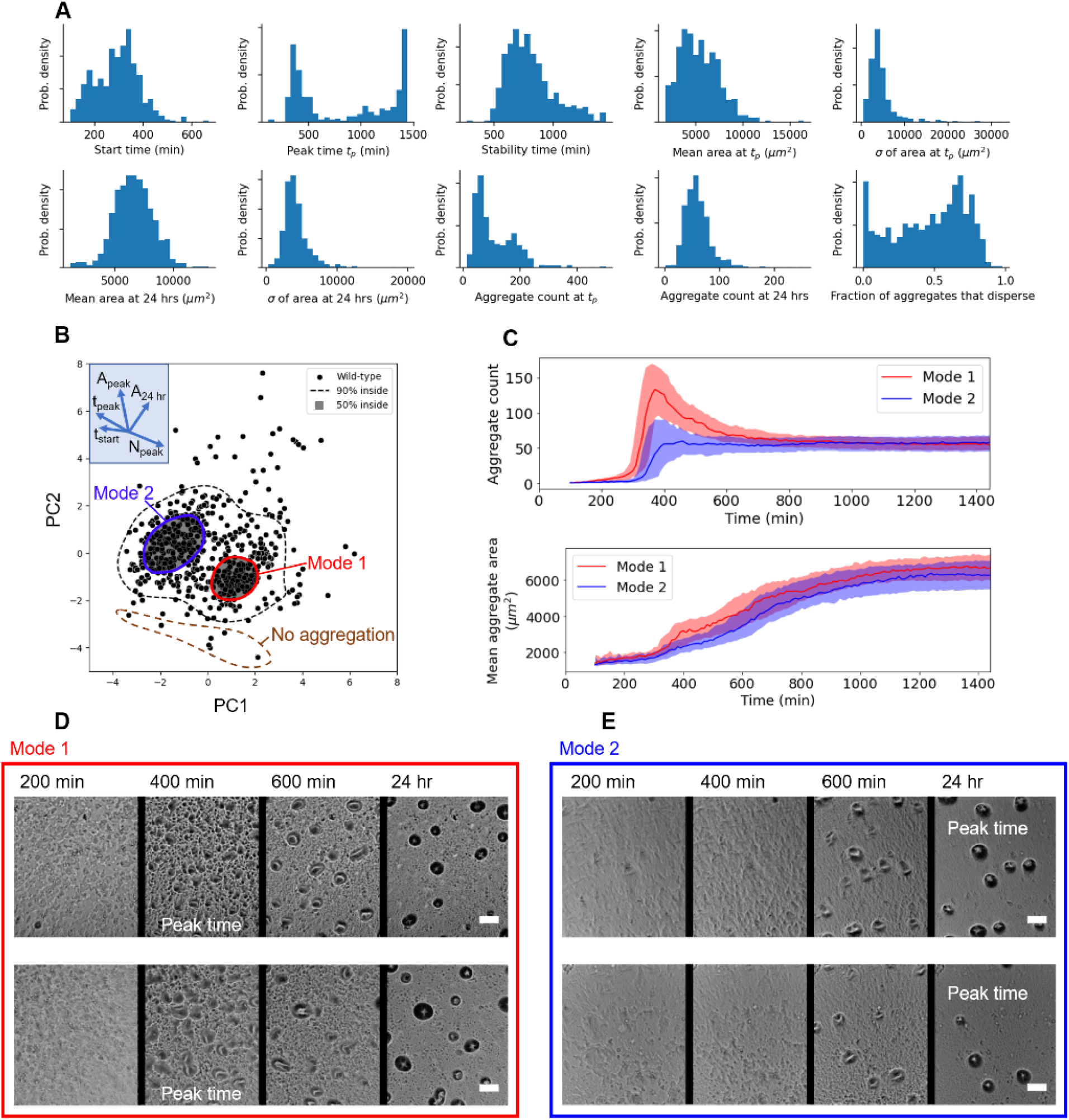
Quantitative breadth of wild-type phenotype. **(A)** Histograms display the range of phenotypic metrics across over 500 wild-type aggregate development time series. Bimodal shapes in peak time (when total aggregate area is maximum), aggregate count at peak time, and fraction of aggregates that disperse reflect the two most common groupings of metrics. Long-tailed distributions, such as standard dev. *ķř*) of area and aggregate count (both at peak time and after 24 hours) indicate the presence of abnormal phenotypes. All y-axes display probability density. **(B)** By using PCA to combine information from all ten metrics, each wild-type time series is plotted as a single datapoint in a phenotypic feature space. PC1 primarily measures aggregate area, and PC2 correlates with number and timing of aggregates. For example, while moving in the direction of the arrow labelled “N_peak_,” datapoints will have generally higher numbers of aggregates at peak time. Units of PC1 and PC2 are arbitrary, although the origin at (0,0) represents average behavior across the full wild-type dataset. A contour is drawn enclosing 90% of the datapoints, separating typical phenotypes from rare phenotypes. Within typical behavior, two separate clusters, Mode 1 and Mode 2, contain 50% of the wild-type datapoints. **(C)** Curves displaying the total number of aggregates over time (top) and mean area of aggregates over time (bottom) illustrate the developmental differences and similarities between the two wild-type modes. The central line represents the median at each time point, and the colored bands span the 25^th^ to 75^th^ percentiles at each time point, i.e. half the data about the median. In Mode 1, a larger number of aggregates develop at an earlier time, most of which disperse. The final number of aggregates is comparable for both modes. The rates of increase of mean area are also similar across the two modes. **(D&E)** Two representative time series each for Mode 1 and Mode 2 phenotypes at three relevant time points. Mode 1 displays many, dense aggregates that form early and then disperse. This causes an early peak time. Mode 2 displays aggregates that form later, most of which persist through the 24 hours of development, slowly growing in area and darkening. This causes a late peak time. Scale bar 100 μm.

We next map the wild-type dataset in a visualizable way. Because we use ten phenotypic metrics, each time series may be represented by a point in a 10-dimensional phenotype space, where points closer together are more phenotypically similar than points far apart. To reduce the number of dimensions but retain the structure of our dataset, we use principal component analysis (PCA), to reduce the number of dimensions from ten to two. The resulting metrics from each time series are mapped to a point in a 2D phenotype space. The two dimensions of this space are called PC1 and PC2, the first and second principal components, respectively. PC1 and PC2 are each a single numerical measure that is a mathematical composite of multiple quantitative features, each weighted differently. Principal component analysis guarantees that PC1 and PC2 are the metrics that display the most variation across the wild-type dataset as compared to any other linearly independent combination of the input metrics. Between just PC1 and PC2, the majority of the variance across the full dataset (56%) is accounted for. The distribution of points in this map constitutes the wild-type phenotype profile.

The unique weighted combination of metrics that make up PC1 and PC2 indicate key metrics that can distinguish behavior. The weights are bounded between −1 and 1, with larger absolute values indicating more strongly weighted metrics. Both PC1 and PC2 contain a mix of all ten metrics, with no one metric standing out in significance over the others, but rather groups of metrics being more significant. In this study, the top weighted metrics of PC1 (with weights given in parentheses) are number of aggregates at peak time (0.47), fraction of aggregates that disperse (0.44), peak time (−0.43), and start time (−0.39). For PC2, the top metrics are mean area at peak time (0.57), standard deviation in area at peak time (0.46), standard deviation in area at 24 hours (0.42), and mean area at 24 hours (0.40). The full list of weights is given in the Supplementary Materials (Table S1). In summary, PC1 is primarily shared between timing and the total number of fruiting bodies that form, in diametrical opposition. That is, when aggregation starts and peaks at an earlier time, the number of fruiting bodies tends to be larger, and vice versa. PC2 is a variable independent from PC1 that mostly characterizes area. Thus, large aggregates can present in large or small numbers, and do so early or late relative to average wild-type behavior.

We observe two primary modes of aggregate formation, as shown by the two shaded regions in Figure 3B. What we term “Mode 1” features aggregates that start forming and peak in total aggregate area generally sooner than other wild-type assays. Mode 1 aggregates are generally numerous, small, and dark at peak time, but a large fraction of them disappear before 24 hours of development. These aggregates tend to be dynamic and lack a well-defined shape until after peak time (Figure 3D). In contrast, Mode 2 aggregation is less mature early on, with either no visible aggregates or aggregates with fewer layers of cells able to block light (Figure 3E). These aggregates are more static and form with more well-defined shapes, and more of them tend to persist through the 24 hours of development. Because these aggregates tend to persist once they form, the time of peak total area is very late for Mode 2, when stable aggregates are still growing slowly. Although there are fewer Mode 2 aggregates at peak time than most wild-type assays, the mean number and size of these aggregates at 24 hours is equal to that of Mode 1, as well as wild-type assays in general. Both modes demonstrate more consistently sized aggregates than other wild-type assays, both at peak time and at 24 hours. Histograms of all ten metrics for the two modes are presented in the Supplementary Materials (Fig S2). Exceptional phenotypes observed in our wild-type dataset include those that produce unusually large fruiting bodies. These occur by a variety of mechanisms, such as large aggregates forming either extremely early with defined shapes from initial formation or extremely late with shapes that only appear visible towards the end of 24 hours (time series in Supplementary Materials, Fig. S3). These abnormal behaviors present at the margins of PCA phenotype space because they represent a confluence of multiple abnormal metrics, revealing more information than standard statistical tests on one metric at a time.

Some rare behaviors observed include failure to aggregate, which occurred in about 2% of wildtype assays, and failure for aggregates to stabilize after 24 hours, which occurred in about 17% of wild-type assays.

We choose four mutant strains to compare with our nominal wild-type strain DK1622, each with 60 to 80 replicates each collected over two to six separate days. These strains were chosen to be developmentally similar to wild-type in order to test the sensitivity of our methods. In preliminary experiments, all four strains produced a set of three replicates that were manually identified as “near-wild-type” and displayed final aggregate size and number that could not be distinguished from wild-type with a Student’s t-test. Three mutant strains contain insertions of simple reporter genes, such as DK10546 producing GFP (See Methods for more details on each strain). Analyzing these strains tests the assumption that introducing reporter genes into a prokaryotic genome will not significantly impact cellular behavior or emergent phenotypes, an important preliminary consideration before their use in other experiments. When more replicates had been analyzed, standard statistical tests distinguish one strain, DK7517, as distinct from wild-type because it produces smaller than average aggregates (Fig. 1). Because the distribution of wild-type final mean areas is non-Gaussian as measured by a Shapiro-Wilk normality test, the Kolmogorov-Smirnov test for distinguishing two distributions was chosen as the standard test in favor of a Student’s t-test, which assumes normality of the underlying distributions.

The developmental data for the additional replicates of the mutant strains are projected onto the same PC1, PC2 axes that were defined for the wild-type data. This allows direct comparison and visualization of multiple metrics simultaneously. The typical behavior and variability of each mutant strain’s development is captured by two regions: a contour is drawn that captures 50% of the datapoints, creating an effective median region in PCA space. We then choose a wider contour that captures 90% of datapoints to serve as a boundary for abnormal phenotypes. By comparing the distribution of the mutant strain points to that of wild-type, a p-value can be calculated for the null hypothesis that the mutant datapoints are drawn from the wild-type distribution. This p-value depends on the number of points found inside the 50% contour and the number inside the 90% contour, and was calculated with bootstrapping, as described in Methods. This is a nonparametric, data-driven statistical method that makes no assumptions about the dataset *a priori*, allowing for multi-modal distributions which are likely to arise in living systems. Validation can be confirmed on multiple subsamples to quantify the impact of day-to-day variation on the p-value.

All four strains demonstrated subtle yet statistically significant departure from wild-type behavior. Strains DK10546 and DK4322 in particular showed a preference for Mode 1 behavior, with Mode 2 being rarely expressed, unlike in wild-type. Some mutant replicates that exhibited rare behaviors are highlighted in Fig. 4B. Replicates of DK10546 displayed more extreme versions of Mode 1 behavior, in which many small aggregates form early on, almost all of which disperse by 24 hours. DK4322 replicates also displayed a more extreme version of Mode 1 behavior in which aggregates at peak time, although distinct, had very irregular shapes. The final aggregates were slightly larger and more varied in area than typical wild-type assays. The aggregates of some DK6665 replicates formed from sparse, small points that formed late and grew steadily over the course of the 24 hours. This nucleation was seldom expressed in wildtype. This strain also displayed difficulty in the dispersal of the random initial cell clumps that are present at inoculation. In wild-type, these initial clumps nearly always disperse, and final aggregate positions have no correlation with these initial clumps. Finally, the abnormal behavior of DK7517 replicates, which involved late aggregates that never significantly darkened, was also a noticeable deviation from even exceptional wild-type behavior. Among the mutant strains, about 2% failed to aggregate (the same fraction as wild-type), and 30% to 45% of mutant assays failed to stabilize after 24 hours, a significant increase from the 17% observed in wildtype.

**Figure 4:**
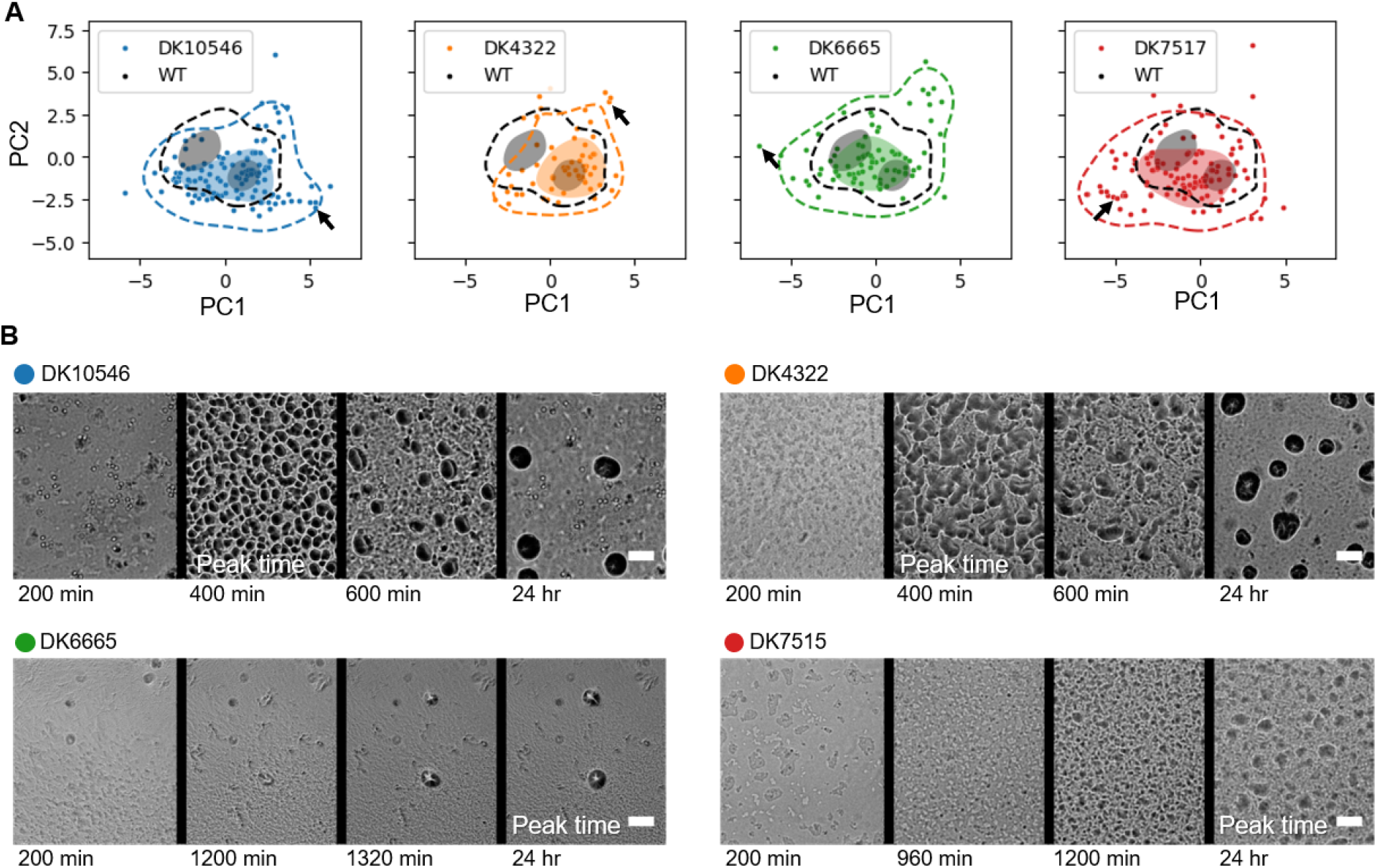
Deviation of near-wild-type mutant strains from wild-type behavior. **(A)** Each mutant development time series is plotted as a single data point in phenotype space, as measured by the collective metrics PC1 and PC2. For each respective strain, dashed contours enclose 90% of data points, and the shaded region(s) enclose 50% of data points. The 90% and 50% contours for wild-type are shown for reference. The deviation of mutant phenotype from wild-type is determined by the departure of the mutant distribution from the wild-type distribution. Statistically significant departures from the wild-type distribution are measured for all four mutant strains, with p-values calculated for subsamples of only 15 replicates each. These p-values are calculated from many random samplings drawn from the wild-type dataset (Methods). Arrows point to time series shown in **(B)** Time series of phenotypes expressed rarely in wild-type are shown at three relevant time points for each mutant strain. Scale bar 100 μm.

## Discussion

Each instance of fruiting body formation is the result of the combined effects of the genetic background, environmental factors both controllable – such as temperature or substrate stiffness – and uncontrollable – such as local pockets of varying initial cell density, or the changes in gene expression that uniquely unfold for that specific population of cells. Genetic changes may be better described by how they affect the odds of a multiplicity of outcomes. In fact, we observe that most mutants behave like wild-type a majority of the time. This is a significant reinterpretation of the meaning of the “phenotype” that results from a given genotype, not as a guaranteed outcome, but as a reshuffling of likely outcomes. This description is appropriate to the physics of living systems, which are tuned through evolution to be poised at the center of a variety of behaviors, ready to adapt to rapid changes either in the organism or its environment. Techniques in the biostatistics community are consistent with this perspective, such as probabilistic latent variable models^11^, which complement the analysis presented in this study.

Essentially, the statistical method reported here characterizes phenotype in terms of abnormal behavior, either locally – i.e. groupings of behavior that fall within the broad scope of wild-type, but are still outside the norm – or globally – i.e. behaviors that are never expressed in the entire wild-type profile. Because measures of mean behavior can fail to capture variations, such as a shifting distribution that happens not to be skewed, the study of abnormalities is a fruitful ground for distinguishing the effects of single-gene mutations, especially when enough replicates can be performed to reliably observe abnormal behavior^8^.

We also quantify the rate at which the wild-type strain fails to create mature fruiting bodies under the conditions given in the Methods section. It is important to distinguish between strains that fail to germinate spores due to a lack of aggregates versus those that aggregate yet still cannot manage to sporulate. It was previously reported that up to 10% of wild-type fruiting body assays cannot successfully germinate new colonies^12^. However, it is unknown to what degree there is a lack of fruiting body morphogenesis during these sporulation assays. The cause for this abnormal phenotype could be disruptions of the signaling pathways necessary for sporulation or a physical issue preventing assembly of aggregates (e.g., low cell density or lack of motility). We show that 2% of our time-lapse images failed to produce aggregates, which would account for only a fraction of the total 10% of instances of failed germination. This does not address what could be happening internally to cells in the remaining instances where fruiting bodies mature with normal dynamics but remain unable to germinate colonies post-starvation. However, there is now a quantification of the failure rate in this system which is crucial to modeling.

Our focus on abnormal development also emphasizes the need to vet exceptional data to ensure that they represent genuine but rare behavior and are not simply abnormal due to a failure of data processing, such as inconsistent imaging conditions. Insofar as possible, the metrics chosen in this study were selected to minimize dependence on imaging setup, and manual vetting was performed on points that fell on the margins of PCA phenotype space.

The methods we report in this work are not unique to bacterial development. A similar analysis could be performed for any stochastic system that can: 1) Have many comparable replicates prepared, 2) Have multiple relevant metrics measured for each replicate. This method can thus be compared to a similar general analysis framework, such as machine learning. Machine learning is powerful in that metrics do not need to be chosen in advance. However, this comes at the cost of the transparency in how categorization is accomplished, and the need for training a model on input data categorized by some other method. It is often true that a system of interest has several obvious aspects that are amenable to measurement with image processing. In the case where the dynamics of the system are relevant, our method is also attractive to use with time series image data, which in raw form can be multiple gigabytes in size and generally difficult to work with on large scales. The initial choice of phenotypic metrics represents a large simplification of the unprocessed image data that ensures phenotypic relevance is preserved over image acquisition noise. The further reduction of the phenotypic dataset from ten to two dimensions not only produces a phenotypic map that is sufficiently navigable to reveal overall structure and guide investigation of individual datapoints, but also avoids a well-known problem in data science associated with the so-called “curse of dimensionality.” This issue occurs in high-dimensional datasets, where geometry tends to make the distance between neighboring points similar to the distance across the dataset, making “similarity” in terms of distance essentially meaningless^13^.

Although 60 replicates or more were analyzed for each mutant strain in this study, a strain that appears like wild-type to the eye can be distinguished from wild-type with fewer replicates. By analyzing the distribution in PCA space of many subsamples of wild-type aggregation, we find that only 15 replicates spread over two days are needed to establish departure from wild-type behavior for each of the mutant strains above. Notably, for this same sample size, standard statistical tests based on individual metrics (average aggregate area after 24 hours shown in Supplementary Materials) can only distinguish one strain, DK7517, as distinct from wild-type, and with less statistical power than the method used in this study. Because the distribution of final mean areas is non-Gaussian as measured by a Shapiro-Wilk normality test, the Kolmogorov-Smirnov test for distinguishing two distributions was chosen as the standard test in favor of a Student’s t-test, which assumes normality of the underlying distributions.

Our results indicate that there are indirect effects on fruiting body formation dynamics in *M. xanthus* due to the use of common reporters. Reporter genes are used as effective markers for successful transfection and are used for quantitative assays, which require them to not obstruct or alter the mechanism of study. GFP, a 28kDa green-fluorescent-protein, allows for the precise visualization of proteins using UV light. Although small enough to diffuse from the cytosol into the nucleus, there are inherent indirect costs to attaching these tags to a molecule of interest. By inserting this extra DNA, another introduction of molecular noise via transcription and translation steps is added during these biochemical reactions^14–16^. These non-target effects can cause cellular differences in expression changes which contribute to the stochastic variation we see in the overall cell population^17^.

Another reporter gene, Tn5 lac, is a promoter-less trp-lac fusion that was designed to identify strains that specifically increase beta-galactosidase expression at some point during *M. xanthus* development as developmental markers. Transposons are a diverse class of mobile genetic elements that can promote genetic rearrangements without a requirement for sequence homology^18^. The Tn5 transposon was inserted so lac Z transcription occurred with exogenous promoters and their promoter strength was quantified^19^ to identify genes that were expressed during *M. xanthus* fruiting body morphogenesis. By attaching to the promoter, it was assumed that lacZ expression would occur in parallel with gene-specific myxospore development without disrupting gene function. However, Tn5 transposon insertions can promote adjacent deletions^20^ which can then disrupt regulatory regions and lead to changes in phenotype from differences in gene expression^21^. The ability to differentiate these transposon insertion strains from wild-type emphasizes the need to assess the impact of reporter genes, especially in biophysical studies that focus on developmental dynamics, where differences may be easier to observe.

These results point to a method of gene annotation that is sufficiently sensitive to identify the impact of single gene mutations that would otherwise be imperceptible. Each mutant strain becomes associated with a signature distribution in a PCA space that can be generally defined and used by any laboratory. Strains with sufficiently similar distributions can then be said to share a function because they impact development in a demonstrably similar way. If the signature distribution of well-understood genes is reported, newly characterized genes of unknown function can be compared to those benchmark distributions. Notably, these signatures are agnostic of any specific biological model, and are based only on visually observable characteristics. Future work can process a library of single-gene knockout strains with unknown function. Over time, this also develops an overall phenome with respect to fruiting body development that is quantitative, and which creates a common language of comparison for the function of many different genes. As more such experiments are done in this framework, either through high-throughput imaging methods like those described here or by the collective efforts of many researchers, developmental phenotypes that are uncommon will be revealed. These unusual events serve to define a boundary on multicellular behavior and can expose the regulatory mechanisms of fruiting body formation when stretched to their limits by stochastic factors alone. These “exceptions” can teach us much about the “rule.”

Among these sources of behavioral change, we expect that small aspects of experimental protocol will have subtle but measurable effects. Over the course of the experiments carried out for this work, a new protocol variable was confirmed that is not normally controlled for in *M. xanthus* culture, namely the age of the agar plate containing colonies to be harvested for liquid bacterial culture. It is expected that reintroducing bacteria to liquid culture will “reset” their metabolic state regardless of what state they were in before, but our analysis revealed preliminary evidence that a colony will “remember” the age of the plate it was harvested from and produce fewer fruiting bodies if the colony grew on the plate for at least three days (Supplementary Materials, Figure S4). Although the mechanism of this memory is unknown, the effect has been consistent, and we expect that other such protocol variables exist that have a measurable phenotypic impact.

The sensitivity of the methods presented here may also be used to measure phenotypic response to changes in a variety of environmental variables. In a similar fashion to running replicates of single-gene mutants, running replicates with differing substrates will reveal subtle or overt changes in phenotype. This future work could address the missing environmental information of the genotype-phenotype problem and expand the bounds of “wild-type behavior” as a function of environmental conditions. The contour bounding abnormal wild-type behavior encompasses exactly what has not yet been characterized to have a specific cause, providing both a measure of ignorance of relevant physical and biological mechanisms, and a way to characterize how much knowledge is gained when subregions can be assigned a root cause, and thus separated from wild-type.

## Conclusions

Overall, our work demonstrates that single gene disruptions can produce measurable changes in *M. xanthus* aggregation development, even when introducing reporter genes widely assumed to be benign with respect to impacting cellular behavior. These effects are subtle and emphasize changes in dynamics over changes only in final developmental outcome, but they can be reliably observed in samples of only 15 replicates. These methods provide a pattern for characterizing development by mapping out phenotype in a clearly visualizable way that also incorporates the many different quantitative aspects that can be measured in collective, living systems. These tools can serve as a language of data presentation with applications in gene annotation or investigations of the impact of environmental variables on the genotypephenotype problem.

## Methods

### Imaging setup

The setup can simultaneously collect time series images for 96 experiments using an array of compact and identical microscopes controlled by a central computer. Each of the 96 microscopes is equipped with a single 4X objective lens, a Peltier device that maintains stage and sample temperature, a red-light source, and a camera controlled by a Raspberry Pi, a single-board minicomputer. The 3D-printed armature and assembly hardware serve to keep all components firmly in place and provides a focus knob for higher image quality.

To ensure uniform control of all microscopes and central storage of their time series output, each individual Raspberry Pi unit is networked via ethernet and two 64-port network switches to a central hub computer. This computer runs Piserver software, which boots each Raspberry Pi from a single operating system image, allowing software to be changed and updated for all Raspberry Pi units simultaneously. Custom software written in Python provides a convenient GUI to control image acquisition from each camera via SSH and organize output in a centralized image storage location.

### Cell culture

Long term stock cultures were recovered on nutrient rich CTTYE media agar (1% Casein Peptone (Remel, San Diego, CA, USA), 0.5% Bacto Yeast Extract (BD Biosciences, Franklin Lakes, NJ, USA), 10 mM Tris (pH 8.0), 1 mM KH(H_2_)PO_4_(pH 7.6), 8 mM MgSO_4_). Cells were harvested from the plates and used to inoculate broth cultures in CTTYE with vigorous shaking at 32°C and grown to an approximate density of 4×10^8^ cells/mL (100 Klett or 0.7 A_550_).

Cells were centrifuged to remove the nutrient broth, washed in TPM buffer (10 mM Tris (pH 7.6), 1 mM KH(H_2_)PO_4_, 8 mM MgSO_4_), and resuspended to a final concentration of 4×10^9^ cells/mL. For the development assay, approximately 4×10^7^ cells (10μL aliquots) were spotted onto a TPM agar slide, a nutrient limited medium, then incubated on the microscope stage at 32°C for 24 hours. TPM slides were prepared as previously described^22^.

The strains used in this study are as follows:

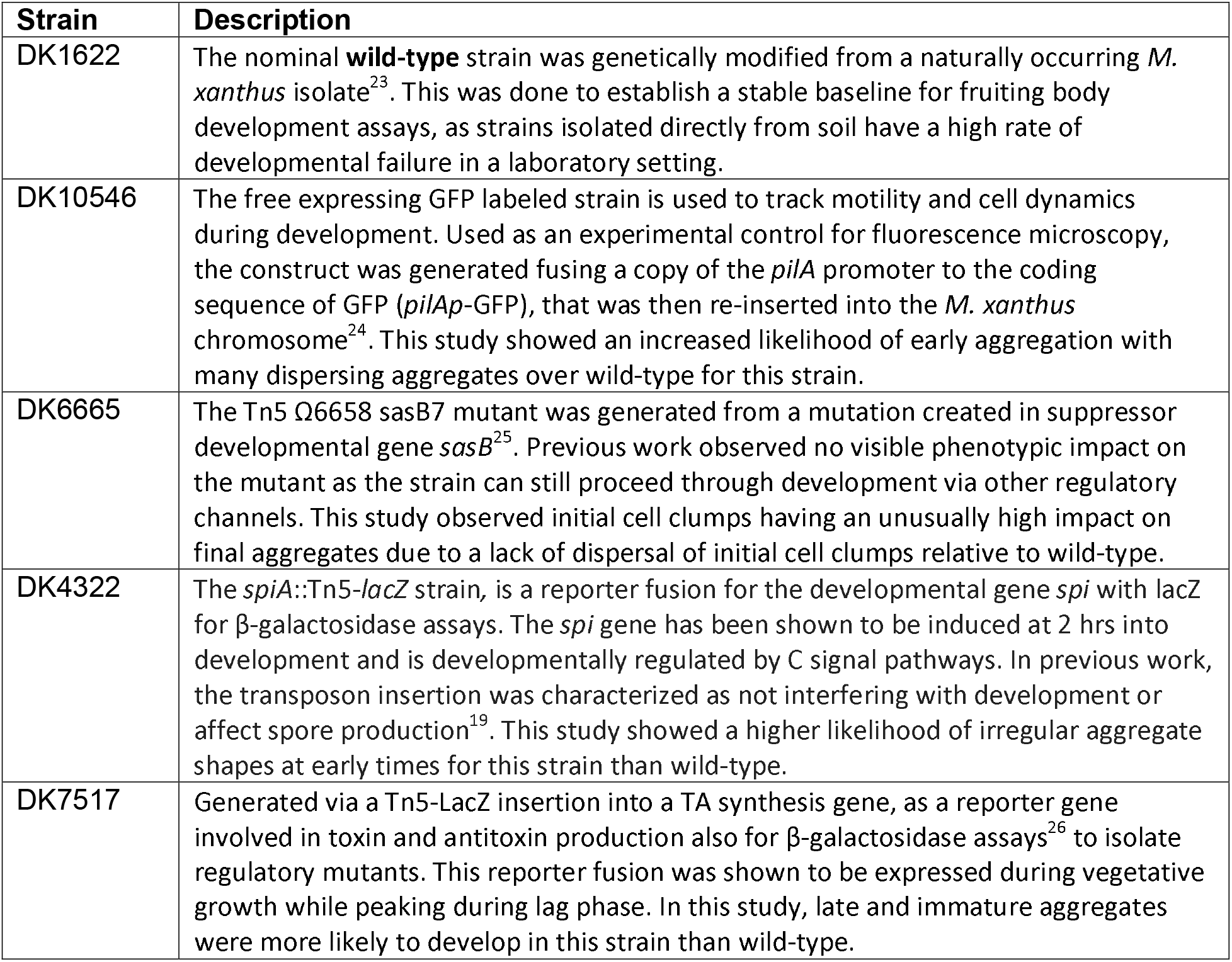

### Image processing pipeline

Phenotype was automatically quantified for each fruiting body aggregation assay in this study by running 144 individual .TIFF images (ten minutes between each frame over 24 hours of total development) from each time series through a custom Python image processing and analysis pipeline to identify in each frame which pixels could belong to a fruiting body, based on their gray value. The information for the position and geometry of each aggregate was filtered to remove noise and spurious aggregates. This detailed data summary for each time series then had a list of ten specific numbers extracted from it, each of which captures one overall feature, such as the time at which aggregation began or the average size of final fruiting bodies. The values of these ten metrics together were then used in further analysis. The full details of the image processing pipeline and all phenotypic metrics are available in the Supplementary Materials Table S2.

### Statistical methods

To calculate p-values that test the null hypothesis of mutant development datapoints in PCA space being drawn from the same distribution as the wild-type development datapoints, we first generate the contours for the wild-type PCA data by starting with Gaussian kernel density estimation (KDE) and using standard root-finding techniques to draw contours from the density estimate that capture 50% and 90% of the PCA datapoints. An appropriate kernel size for the KDE is validated by using 75% of the wild-type dataset, and ensuring that, across many subsamples of the remaining 25% (verification data), the distribution of enclosed points is centered on the appropriate percentage. When this distribution is skewed, it indicates overfitting of the original contour. With these contours drawn, we then use a data-driven statistical technique similar to bootstrapping. Given a sample size N, 10,000 samples of that size are drawn from the wild-type dataset. Each subsample has a characteristic pair of numbers, (*n_50_*, *n_90_*), which corresponds to the number of points in the sample that fall inside the 50% and 90% contours, respectively. Once the distribution of these pairs for wild-type data is known, *n_50_* and *n_90_* are calculated for a sample of mutant PCA datapoints of size N. The fraction of wild-type videos that have both *n_50_* and *n_90_* greater than the mutant sample’s values of *n_50_* and *n_90_* gives the p-value, or the probability that a sample of wild-type data of size N would exhibit the same distribution. Contours for mutant strains are shown for visualization only, and do not figure into the calculation of the p-values.

## Acknowledgements

The authors acknowledge funding from NSF MCB 2026747 and NSF DEB 2033942 awarded to A.P. and NSF MCB 1856665 and NSF DMS-NIGMS 1903160 awarded to R.W.

