## Supplemental Information for "Stochastic bounds of aggregation dynamics distinguish near-wild-type from wild-type strains in social bacteria"

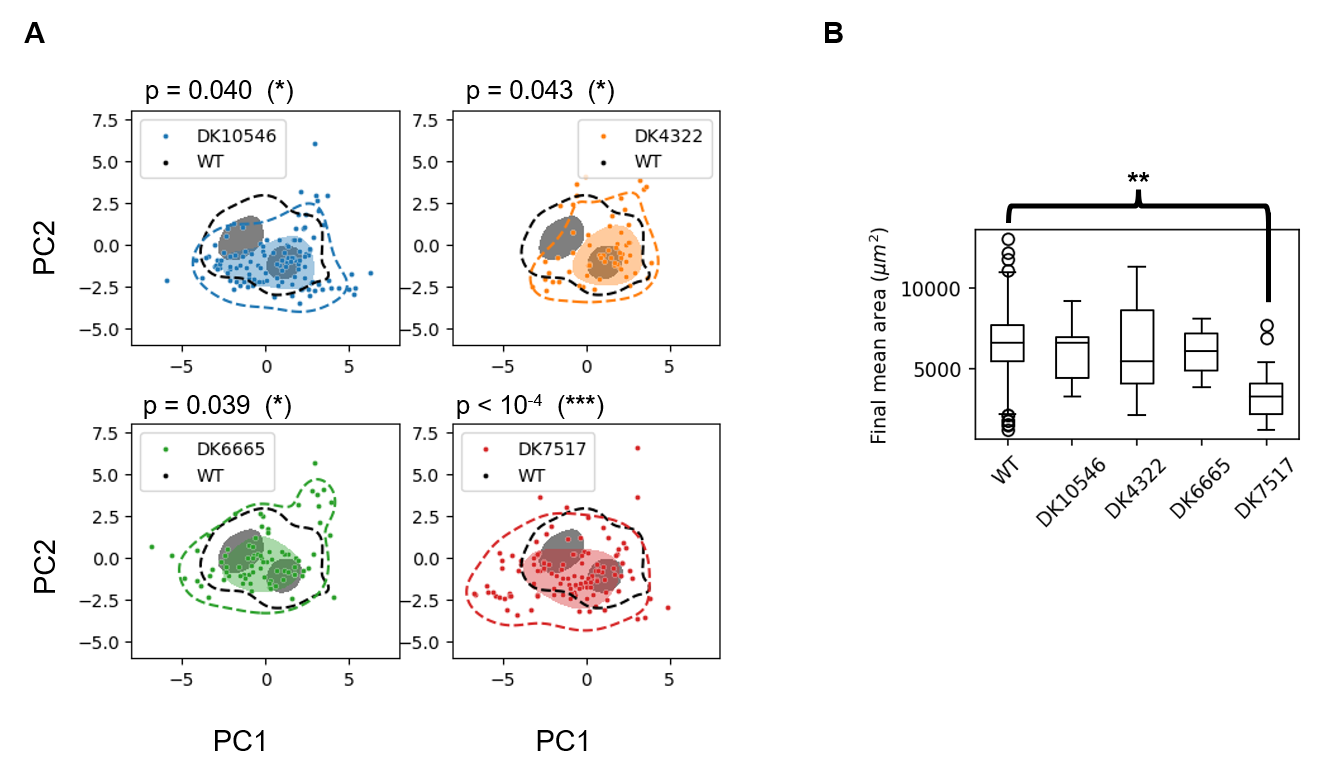

**Figure S1: Standard statistical test (Kolmogorov-Smirnov on one metric) compared with p-value calculated from changing distributions in PCA space.** **(A)** Each mutant development time series is plotted as a single data point in phenotype space, as measured by the collective metrics PC1 and PC2. For each respective strain, dashed contours enclose 90% of data points, and the shaded region(s) enclose 50% of data points. The 90% and 50% contours for wild-type is shown for reference. The deviation of mutant phenotype from wild-type is determined by the departure of the mutant distribution from the wild-type distribution. Statistically significant departures from the wild-type distribution are measured for all four mutant strains, with p-values calculated for subsamples of only 15 replicates each. These p-values are calculated from many random samplings drawn from the wild-type dataset. **(B)** The same mutant strains are compared against wild-type using only a metric taken from the last frame of development, final aggregate area. With subsamples of size N=15, a Kolmogorov-Smirnov test can only distinguish between DK7517 and wild-type with statistical significance.

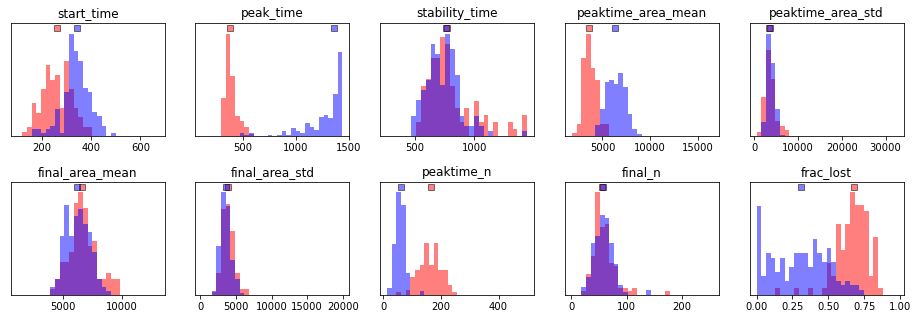

**Figure S2: Comparison of metrics for Mode 1 (red) and Mode 2 (blue) wild-type assays.** Modes 1 and 2 are defined by the two regions of the contour bounding 50% of the total wild-type assay data in PCA space, as shown in Fig. 3 of the main text. The medians of each histogram are indicated by the position of the square marker at the top of each subplot.

A fruiting body aggregation assay performed in any lab can be compared to the results reported in this work by calculating the values of PC1 and PC2 for that assay.

First, scale each metric by subtracting the mean WT value of that metric and dividing the result by the WT standard deviation of that metric, according to the values reported in Table 1. This normalizes each metric to have zero mean and unit variance. Then, each resulting scaled metric is multiplied by an appropriate weight, either for PC1 or PC2. The weighted sum of the scaled metrics gives the final value of PC1 or PC2 for that assay.

| **Metric Name** | **Mean value (WT dataset)** | **Standard dev. (WT dataset)** | **Weight (PC1)** | **Weight (PC2)** |
| --- | --- | --- | --- | --- |
| Start time (min) | 291 | 90.0 | -0.387 | 0.047 |
| Peak time (min) | 843 | 454 | -0.430 | 0.245 |
| Stability time (min) | 1008 | 486 | 0.183 | 0.038 |
| Mean area at peak time (μm^2^) | 5414 | 2063 | -0.126 | 0.574 |
| Std area at peak time (μm^2^) | 4670 | 3525 | 0.188 | 0.465 |
| Final mean area (μm^2^) | 6617 | 1690 | 0.255 | 0.401 |
| Final std area (μm^2^) | 4244 | 1972 | 0.281 | 0.424 |
| N at peak time* | 113 | 75.6 | 0.468 | -0.198 |
| Final N* | 61.7 | 27.8 | 0.149 | 0.062 |
| Fraction lost | 0.463 | 0.253 | 0.443 | -0.083 |

**Table S1: Numerical definition of PC1 and PC2 for reproducibility.**
*Number of aggregates is reported in a field size of area 5.0 mm^2^. Different field sizes should scale aggregate count appropriately, assuming a constant density of aggregates per mm^2^.

| Metric Name | Description | Formula |
| --- | --- | --- |
| Start time | The time elapsed between inoculation and the beginning of observable aggregation. | The earliest time at which at least ten aggregates have reached an area of at least 800 μm^2^ |
| Peak time | The time elapsed between inoculation and the moment aggregation reaches maximum total area | The time at which the sum of the areas of all aggregates in one frame is at a maximum value across the time series |
| Stability time | The time elapsed between inoculation and the moment the number of aggregates becomes stable | If stability is achieved within 24 hours, the time at which the rate of change of number of aggregates falls below and stays below 0.5 per minute. Rate of change is calculated with a Savitsky-Golay filter using a window 31 minutes wide. Otherwise, 2000 minutes, to represent a later eventual stability time. |
| Mean area at peak time | The average area of all aggregates at the moment of peak time | Peak time is evaluated, and the average area of each aggregate in that one frame is calculated |
| Std area at peak time | The standard deviation in the area of all aggregates at the moment of peak time | Peak time is evaluated, and the sample standard deviation of each aggregate area in that one frame is calculated |
| Final mean area | The average area of all aggregates 24 hours after inoculation | For the one frame showing 24 hours after inoculation, the average area of each aggregate is calculated |
| Final std area | The standard deviation in the area of all aggregates 24 hours after inoculation | For the one frame showing 24 hours after inoculation, the sample standard deviation of each aggregate area is calculated |
| N at peak time | The number of aggregates at peak time | Peak time is evaluated, and all aggregates are counted in the 5.0 mm^2^ visible field in that one frame |
| Final N | The number of aggregates 24 hours after inoculation | All aggregates in the 5.0 mm^2^ visible field are counted in the one frame showing 24 hours after inoculation |
| Fraction lost | The fractional change in number of aggregates from the moment of maximum number to 24 hours after inoculation. | For N_max the maximum number of aggregates (smoothed by averaging the three largest values) and N_24 the number of aggregates 24 hours after inoculation, (N_max – N_final) / N_max |

**Table S2: Description of each of the ten developmental metrics used to quantify aggregation phenotype.**

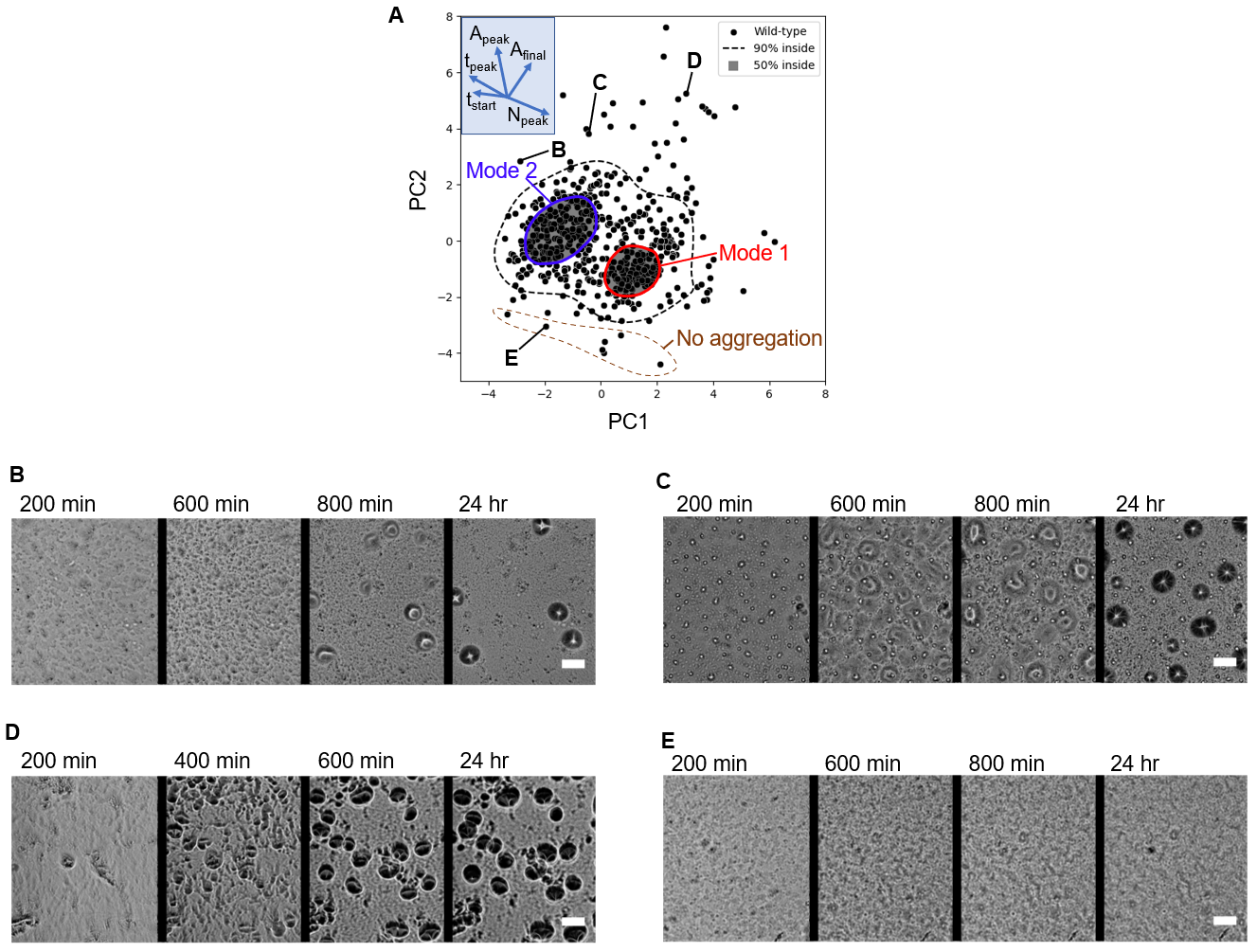

**Figure S3: Abnormal phenotypes expressed in wild-type. (A)** Wild-type phenotype profile, showing the location in PCA space of four instances of illustrative, abnormal behavior in wild-type replicates. **(B)** An extreme instance of Mode 2 behavior, especially due to late start time and the small number of final aggregates. Successful aggregates that are less mature than these are expressed seldom in wild-type. **(C&D)** Time series of abnormal behavior show the larger aggregates that form in the region with high PC2 values, but with significant differences in timing and number depending on the value of PC1. (**E)** A representative time series of failed aggregate formation. Bacterial activity is visible, but aggregates of any significant size or darkness do not form after 24 hours. Scale bar 100 μm.

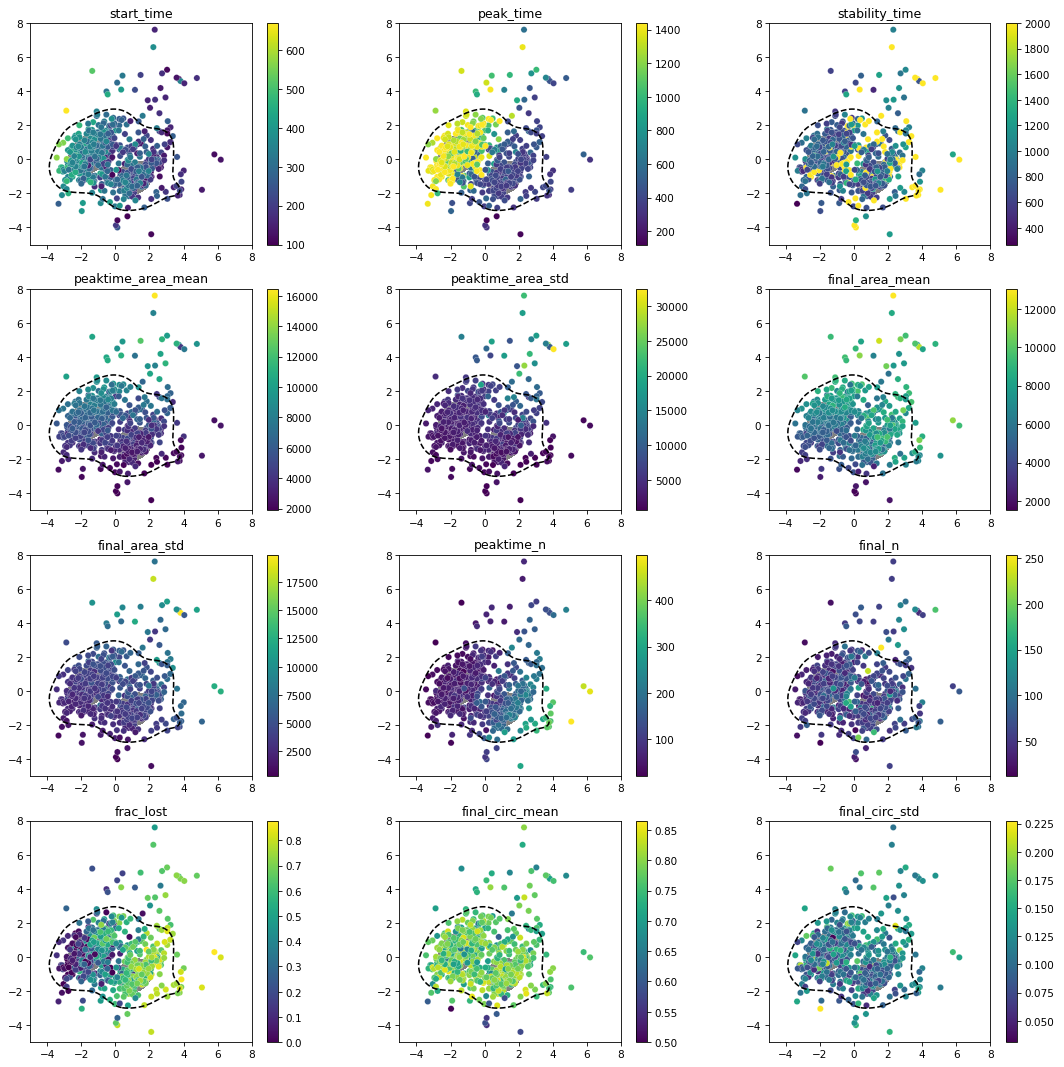

**Figure S4: Variation of metrics across wild-type dataset.** This figure illustrates how PC1 and PC2 organize the variation of phenotypic metrics in PCA space. Of the ten metrics used in the PCA, eight display a particular gradient direction. Only two, stability time and final number of aggregates, do not display a clear gradient direction but vary in a more complex pattern. For instance, high stability time (i.e. those time series that do not stabilize by 24 hours) and high final number of aggregates mostly present outside the regions of Mode 1 and Mode 2 (See Fig. 3 in the main text). Although not included in the PCA, the mean and standard deviation of aggregate circularity at 24 hours is also reported, not showing any clear correlation with the other metrics.

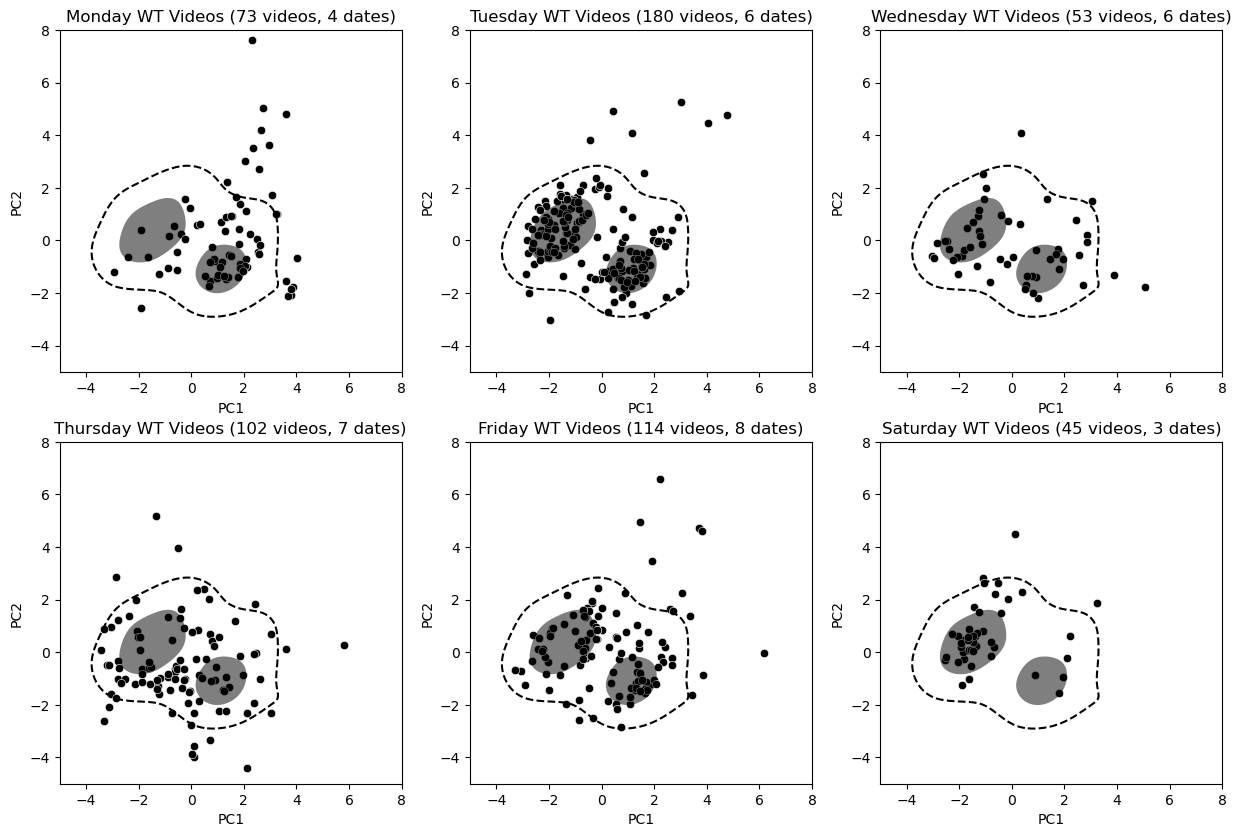

**Figure S5: Variation of wild-type videos based on date of inoculation.** Differences were observed in the distribution of wild-type outcomes depending on the day the time lapse was started. This may represent an effect of the length of time *M. xanthus* bacteria spend in a colony on agar before being transferred to liquid culture and eventually inoculated on the non-nutritive agar used in fruiting body development assays. A shift from Mode 1 to Mode 2 being favored occurs when comparing Monday to Saturday development videos.
